# 3D structure and *in situ* arrangements of CatSper channel in the sperm flagellum

**DOI:** 10.1101/2021.06.19.448910

**Authors:** Yanhe Zhao, Huafeng Wang, Caroline Wiesehoefer, Naman B. Shah, Evan Reetz, Jae Yeon Hwang, Xiaofang Huang, Polina V. Lishko, Karen M. Davies, Gunther Wennemuth, Daniela Nicastro, Jean-Ju Chung

## Abstract

The sperm calcium channel CatSper plays a central role in successful fertilization as a primary Ca^2+^ gateway into the sperm flagellum. However, CatSper’s complex subunit composition has impeded its reconstitution *in vitro* and structural elucidation. Here, we applied cryo-electron tomography to visualize the macromolecular organization of the native CatSper channel complex in intact mammalian sperm, as well as identified three additional CatSper-associated proteins. The repeating CatSper units form long zigzag-rows in four nanodomains along the flagella. In both mouse and human sperm, each CatSper repeat consists of a tetrameric pore complex. Murine CatSper contains an additional outwardly directed wing-structure connected to the tetrameric channel. The majority of the extracellular domains form a canopy above each pore-forming channel that interconnects to a zigzag-shaped roof. The intracellular domains link two neighboring channel complexes to a diagonal array. The loss of this intracellular link in *Efcab9*^-/-^ sperm distorts the longitudinally aligned zigzag pattern and compromises flagellar movement. This work offers unique insights into the mechanisms underlying the assembly and transport of the CatSper complex to generate the nanodomains and provides a long-sought structural basis for understanding CatSper function in the regulation of sperm motility.

---

Freshly ejaculated mammalian sperm must undergo a physiological process called capacitation to be capable of fertilizing the egg^1,2^. The crucial change that occurs during capacitation represents a motility change, i.e. the sperm flagellum beats vigorously and asymmetrically, producing a whip-like motion. This motility pattern – known as hyperactivated motility – enables the sperm to reach the egg by overcoming the viscous microenvironment of the female reproductive tract. Additionally, hyperactivation allows sperm to push through a sticky egg coat, and eventually fertilize the egg^3^. Hyperactivation is triggered by the elevation of the intraflagellar calcium that requires the sperm-specific and Ca^2+^-selective CatSper channel^4,5^. CatSper loss-of-function abrogates hyperactivation of the sperm flagellum and renders males infertile in both mice and humans^6^.

Previous studies have found that CatSper is the most complex ion channel known, with at least ten proteins: four subunits that form a heterotetrameric channel (CATSPER1-4)^5,7^, as well as six additional, non-pore forming subunits, including four transmembrane (TM) proteins with large extracellular domains (ECD) (CATSPERβ, γ, δ, and ε)^8-11^ and two smaller cytoplasmic, calmodulin (CaM)-IQ domain proteins that form the EFCAB9-CATSPERζ complex^8,12,13^ (Extended data Table 1). Deletions or mutations of any of the pore-forming or other TM-subunits results in the loss of the entire CatSper channel complex^6^. Super resolution light microscopy showed that the CatSper channel complex is restricted to four linear compartments within the flagellar membrane^14,15^, generating a unique longitudinal signaling nanodomain in each flagellar quadrant. Genetic evidence suggested that this high-order arrangement is essential for Ca^2+^ signaling and sperm hyperactivation, highlighting physiological relevance of the spatial organization. Disrupting the integrity of the linear nanodomains alters the flagellar waveform and prevents sperm from efficiently migrating *in vivo*^8,14,16^. Specifically, the absence of the cytoplasmic EFCAB9-CATSPERζ complex in *Efcab9*^-/-^ and/or *Catsperz*^-/-^ mutant sperm alters the continuity of each CatSper nanodomain^8,12^, suggesting a regularly repeating, quaternary structure of the CatSper complex within the nanodomains.

Despite many important discoveries mentioned above, the fundamental structure of the native channel complex and its molecular architectural arrangement were still not known. Here, we address these questions by visualizing in-cell organization and domain structures of the CatSper channel complex in intact mouse and human sperm flagella using cryo-electron tomography (cryo-ET).

## Macromolecular composition of the CatSper complex

Ten components have been validated to comprise the CatSper channel complex in the linear nanodomains^6^. However, we previously showed by comparative mass-spectrometry that in mouse *Catsper1*^-/-^ sperm four additional proteins were reduced: C2CD6 (<u>C2 C</u>alcium-dependent <u>D</u>omain-containing protein <u>6,</u> also known as ALS2CR11), E3 ubiquitin-protein ligase TRIM69, SLCO6C1 (<u>s</u>olute <u>c</u>arrier <u>o</u>rganic anion transporter family, member 6c1), and the DNA-binding ATPase FANCM (<u>F</u>anconi <u>an</u>emia, <u>c</u>omplementation group <u>M</u>)^12^ (see also Extended Data Fig. 1a).

To test whether these four candidates are truly associated with the CatSper channel, we used western blot analyses of *Catsper1*^-/-^ and *Catsperd*^-/-^ sperm that lack the entire CatSper channel complex^9,12,14^, and found that the protein levels of C2CD6, TRIM69, and SLCO6C1, but not FANCM, were indeed reduced (Fig. 1a and Extended Data Fig. 1b, c). Moreover, using fluorescence and 3D structured illumination microscopy (3D SIM) we showed that the three CATSPER1/δ-dependent proteins, *i*.*e*. C2CD6, TRIM69, and SLCO6C1, localize in the principal piece - the longest part of the sperm flagellum harboring the CatSper channel - and display the same quadrilinear distribution with the nanodomains (Fig. 1b, c, and Extended Data Fig. 1c-f). In the absence of EFCAB9 and/or ζ, the continuous distribution is disrupted (Fig. 1c and Extended Data Fig. 1f, *lower*), resembling previously reported results for the known CatSper subunits^8,12^. This typical dependence of protein levels and localizations on other CatSper components suggest that they are likely three new *bona fide* CatSper-associated proteins of murine sperm. In particular, it is intriguing to find an organic anion transporter, SLCO6C1 that is highly expressed in rodent testis^12,17,18^, in complex with an ion channel, the murine CatSper. Whereas the loss-of-function effect of C2CD6 remains to be determined, *Trim69*^19^ and *Slco6c1*^20^ are not essential for fertility, indicating their function on the CatSper channel is likely to be modulatory and/or indirect.

**Fig. 1.**
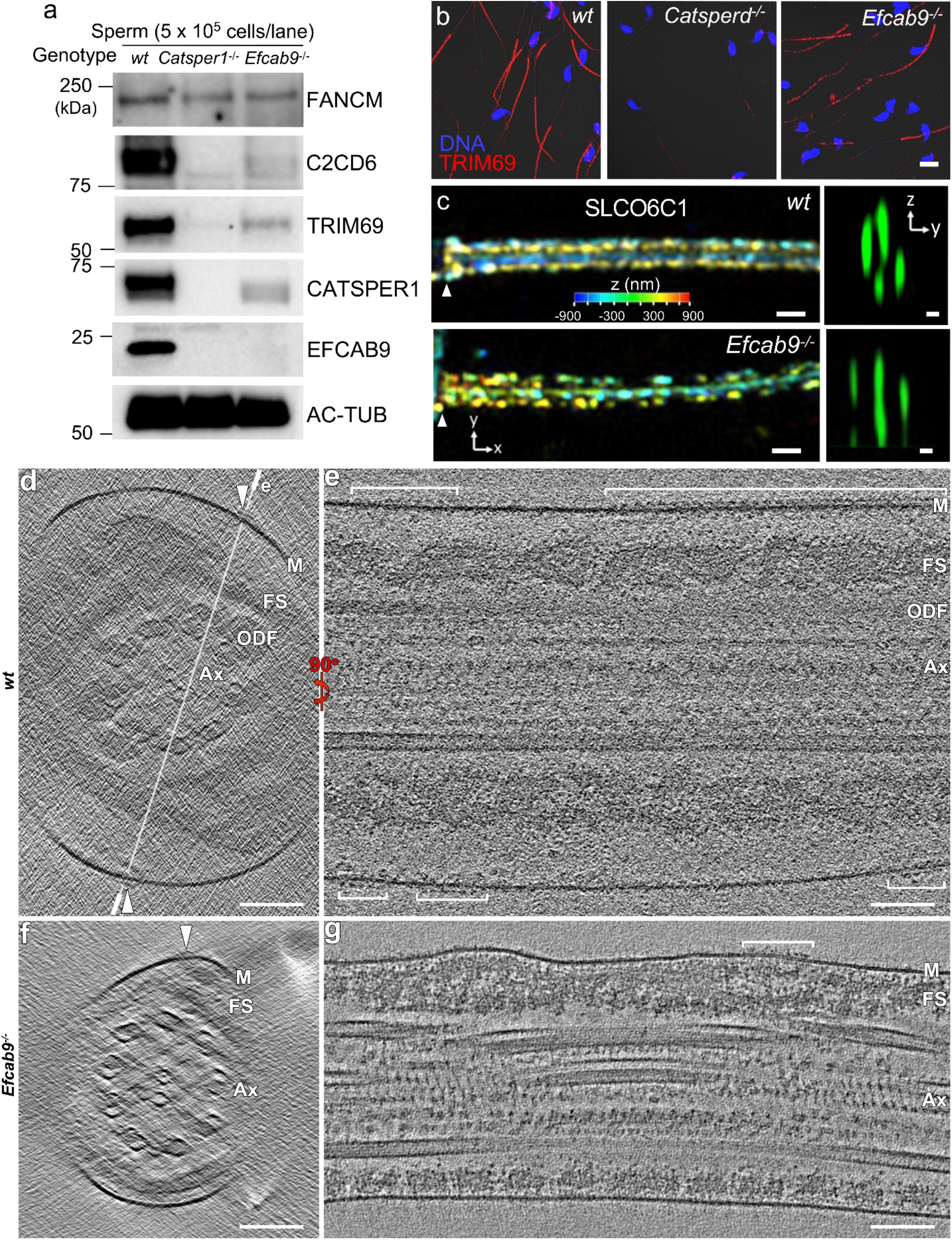
New CatSper components and cryo-ET of mouse sperm flagella visualizing particles of CatSper complexes. **a**. Western blot analyses of four candidate CatSper-associated proteins in the whole cell proteome of mouse wild type, *Catsper1*^-/-^ *and Efcab9*^-/-^ sperm. **b-c**. Immunolocalization of TRIM69 (**b**, confocal light microscopy; sperm head stained blue using Hoechst) and SLCO6C1 (**c**, 3D SIM) in sperm from wild type, *Catsper1*^-/-^, and/or *Efcab9*^-/-^ mice. In (**c**) colors in *xy* projection encode the relative distance from the focal plane along the *z* axis. Arrowheads in each panel indicate the annulus, the junction between the midpiece and principal piece of the sperm tail. *y*-*z* cross sections are shown on the right. Scale bar, 10 μm in **b**; 500 nm in **c** (left); 200 nm in **c** (right). **d-g**. Tomographic slices of representative principal piece regions of mouse sperm flagella show CatSper complexes (arrowheads) viewed in cross section (left) and longitudinal section (right): intact wild type (**d, e**) and *Efcab9*^-/-^ (**f, g**) sperm in non-capacitated state. Other labels: M, membrane; ODF, outer dense fiber; FS, fibrous sheath; Ax, axoneme; CP, central pair. Scale barx, and the recently reported 1:1 stoichiometry of seven TM CatSper subunits in sea urchin sperm (*i*.*e*. CATSPER1-4, β, γ, and ε)^21^, we hypothesize that a single mouse or human CatSper complex might form a nearly half-megadalton extracellular domain (ECD) (Extended Data Table 1), a size that could be visualized by cellular cryo-ET. Therefore, we performed cryo-ET on intact murine and human sperm flagella to characterize the native CatSper complex *in situ*, which avoids potential purification artefacts. Viewing the 3D reconstructed sperm and flagellar membranes in cross-section, we observed protruding particles of ∼25 nm in width positioned to either side of the longitudinal column of the fibrous sheath in the principal piece (Fig. 1d), consistent with the localization for CatSper nanodomains as seen by immuno-electron microscopy (EM)^14^. Out of the four quadrants, only up to two could be visualized in the cryo-tomograms due to the missing wedge effect that results from a limited tilt-angle range in single-axis ET. Longitudinal tomographic slices of the wild type sperm flagella revealed long continuous rows of densely packed particles with an apparent periodicity of 17.6 nm (Fig. 1e). The resolution of reconstructed whole murine flagella was limited due to the ∼900 nm sample thickness of the proximal region of the principal piece. Therefore, we also used cryo-FIB milling to generate ∼200 nm thick slices (called “lamella”) of murine sperm flagella (Extended Data Fig. 2a-g) that resulted in higher-resolution tomographic reconstructions (Extended Data Fig. 2h, i; Extended Data Table 2).

Due to all-or-none assembly of CatSper TM subunits in mouse sperm, knockout of any one of these TM subunits leads to mutant sperm that do not form the nanodomains as they lack the entire CatSper complex^6^ (see also Extended Data Fig. 1a, b; *Catsper1*^-/-^ and *Catsperd*^-/-^). By contrast, *Catsperz*^-/-^ and/or *Efcab9*^-/-^ sperm assemble the CatSper complex missing only the two interdependent non-TM EFCAB9 and CATSPERζ subunits^12^ (see also Fig. 1a and Extended Data Fig. 1b). Because previous observations by super resolution light microscopy and scanning EM suggested the linearity of the nanodomains is discontinuous in a fragmented pattern in *Catsperz*^-/-^ and *Efcab9*^-/-^ sperm^8,12^, we next looked at *Efcab9*^-/-^ sperm for CatSper particles. Indeed, we observed that the particles positioned to the corresponding locations in the flagellar membrane of *Efcab9*^-/-^ sperm formed discontinuous rows, short clusters or individual repeat units (Fig. 1f, g). Together with the position of these particles along flagella, this genetic evidence, *i*.*e*. disruption of the particle-rows in *Efcab9*^-/-^ sperm, strongly supports that these particles are macromolecular CatSper channel complexes that form the quadrilinear nanodomains.

## Zigzag arrangement of CatSper complexes

Slicing the rows of CatSper channel complexes in longitudinal orientation parallel to the flagellar membrane (top-down view) revealed continuous rows with repeating units in a zigzag arrangement of ∼25 nm in width (Fig. 2a-h), demonstrating the CatSper complexes are repeated within the rows. We found that the number of zigzag rows per nanodomain varies from a single row (Fig. 2a, b), two rows that can be up to 100 nm apart (Fig. 2c, d and Extended Data Video 1) or merge into one row (Fig. 2e, f), or up to as many as five parallel rows (Fig. 2g, h). In tomograms of *Efcab9*^-/-^ sperm flagella, we observed mostly short clusters containing only 1-7 units (Fig. 2k-n) and very few continuous rows with a maximum of ∼70 repeats (Fig. 2i, j and Extended Data Fig. 3a, b). Interestingly, the short mutant clusters are no longer well-aligned with the flagellar axis and adopt various angles - up to almost perpendicular – relative to the longitudinal axis of the flagellum (Fig. 2l, n). In *Efcab9*^-/-^ sperm, C2CD6, TRIM69, and SLCO6C1 proteins were reduced but detectable (Fig. 1a, b and Extended Data Fig.1b) as seen for all the previously known 8 TM subunits^12^. These results suggest that the absence of the EFCAB9-CATSPERζ complex from the intracellular side of the channel disrupts the high-order arrangement of the CatSper channel complex and the linear alignment in the nanodomains. Cryo-tomograms of human sperm flagella revealed similar linear rows that are ∼24 nm wide and consist of repeating units that are also arranged in a zigzag pattern (Fig. 2o-r).

**Fig. 2.**
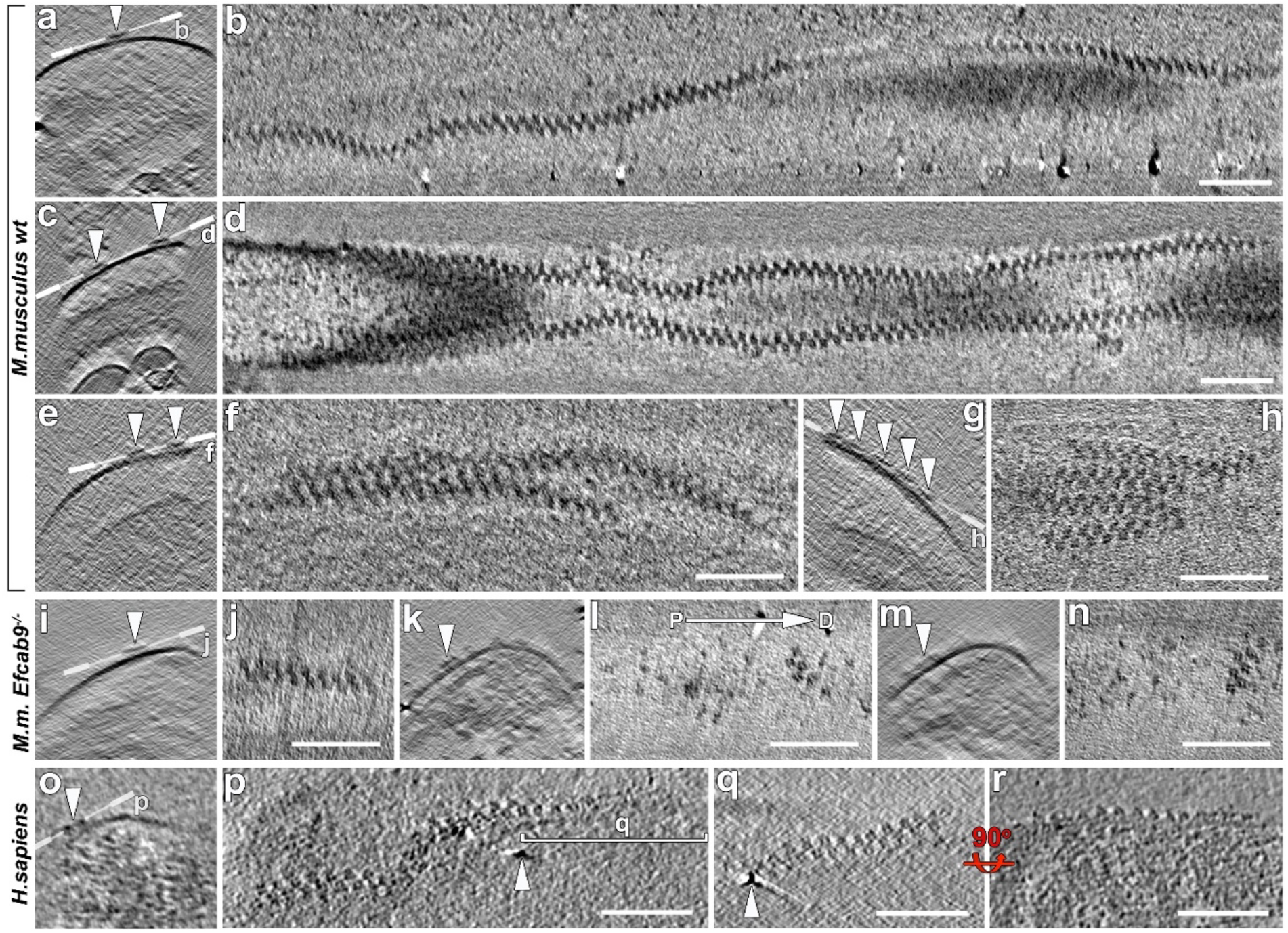
In-cell structure of the native CatSper complexes in intact sperm flagella. **a-h**. Representative tomographic slices of the repeating CatSper channel complexes arranged as zigzag-rows along the longitudinal axis of wild type flagella (cross view from proximal to distal: **a, c, e, g**; top-down view with proximal side of the flagellum on the left: **b, d, f, h**). The number of zigzag-rows (arrowheads) varied from a single row (**a, b**), two rows (**c, d**), merging rows (**e, f**), to up to five rows (**g, h**). **i**-**n**. Representative tomographic slices of *Efacb9*^-/-^ sperm show fragmented, short CatSper complex clusters with altered orientation relative to the flagellar axis (cross section view: **i, k, m**; top-down view: **j, l, n**, the direction from proximal (P) to distal (D) as indicated). **o-r**. Zigzag-arrangement of CatSper in human sperm flagellum (cross section view: **o**; top-down view: **p, q**; side view, **r**). Scale bar, 100 nm.

## Extracellular structures of CatSper form canopy tents that connect pore-forming channels as beads on a zigzag string

After determining the periodicity of the CatSper complexes within the zigzag rows, we performed subtomogram averaging of the repeating units to increase the signal-to-noise ratio and thus the resolution. We averaged ∼2500 CatSper complex repeat units (which includes the application of two-fold symmetry) from continuous rows from 11 acquired cryo-electron tomograms of both whole cells and cryo-FIB milled mouse wild type flagella (Extended Data Table 2). The averages depict unprecedented details of CatSper complexes *in situ* (Fig.3a-h) with up to 22 Å resolution (0.5 FSC criterion; Extended Data Fig.3h, Extended Data Table 2).

**Fig. 3.**
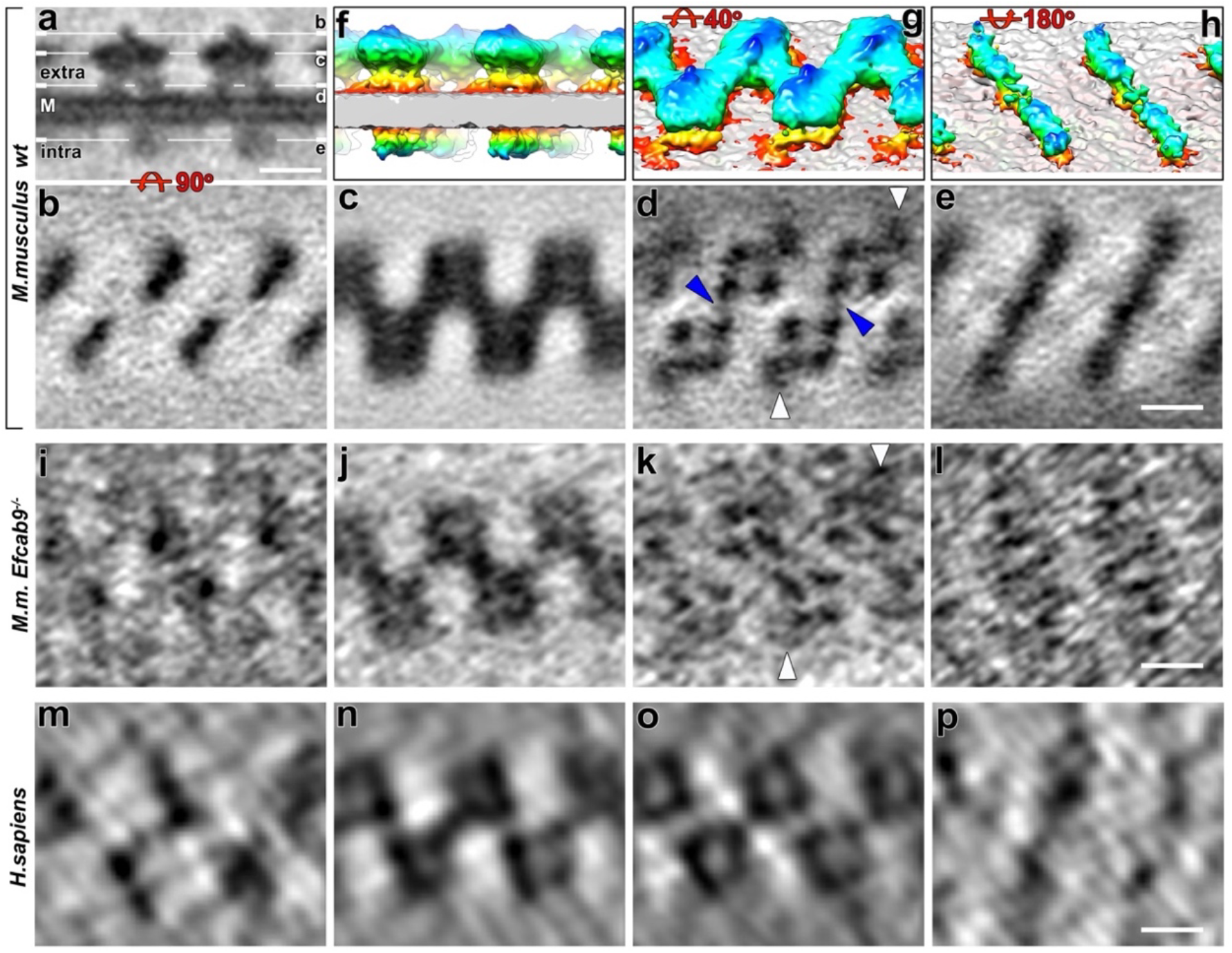
Structural features of CatSper complex in three dimensions. **a-e**. Tomographic slices show the averaged CatSper complex structure from wild type mouse sperm in side (**a**) and top views (**b-e**). **f-h**. 3D isosurface renderings of the averaged CatSper complexes in wild type mouse sperm: **f**, side view; **g**, extracellular domain; **h**, intracellular domain. **i-l**. Tomographic slices show the averaged CatSper complex structure in *Efacb9*^-/-^ sperm in a backslash direction. **m-p**. Tomographic slices show the averaged CatSper complex structure in human sperm flagellum. Lines in (**a**) indicate the slice positions showing the following structural features: **b, i, m**, roof ridge; **c, j, n**, canopy roof; **d, k, o**, tetrameric channel; **e, l, p**, channel intracellular domain. Other labels: white arrowheads, wing structure; blue arrowheads, inner connection between channel subunits, M, membrane. Scale bar, 10 nm.

As shown in Figure 3 and Extended Data Video 2, the averaged 3D structure of the zigzag row reveals that the CatSper complexes are evenly spaced in two anti-parallel lines, *i*.*e*. the complexes are 180º rotated between the two lines. The appearance of an ∼25-nm-wide zigzag-pattern results from the staggering of the rows of channels and the ECDs connecting across the lines (Fig. 3c, d, g). Several structural features of the whole channel unit are visualized from extra-to intracellular domains across the inner and outer leaflet of the membrane bilayer (Fig. 3a-h). In the side view (Fig. 3a), the most prominent structural feature of each CatSper complex is the uniquely shaped ECDs that form a 11.2 nm high canopy tent in which the majority of the ECD mass forms the canopy roof (Fig. 3a, f, g). The roof is connected between neighboring complexes to a continuous zigzag ribbon around 6.6 nm away from the flagellar membrane (Fig. 3c, g).

Tangential slices (*i*.*e*., top views) through this extracellular part, show closest to the membrane clearly the asymmetric unit: the tetrameric arrangement of the CATSPER1-4 subunits with an additional density that we named “wing” at the outside corners (Fig. 3d, white arrowheads). The position of the wing clearly reveals the 180º rotation between connected neighboring CatSper complexes. At an inside corner of the tetramer – opposite to the wing-connected subunit – at least one of the pore-forming channel subunits forms a fine but clearly visible connection to the identical subunit of the “forward slash”-neighboring, 180º rotated CatSper channel (Fig. 3d, blue arrowheads). The diameter of the tetrameric channel is 10.6 nm, which is in a similar range with the size observed for other tetrameric channels such as 10 nm wide Cav1.1^22^. The center-to-center spacing between channels along the zigzag string is 15 nm.

The ECD canopy roof is positioned right above the tetrameric channel (*i*.*e*., the four tent poles) (Fig. 3a-d). Interestingly, the roof ridge (Fig. 3b; Fig. 3f-g, dark blue) is off-center and tilted in the same “forward slash” direction as the two connected inner pore-forming subunits (Fig. 3b, d). Based on the subtomogram average, the mass estimation of the ECDs of one CatSper channel complex is ∼430 kDa, close to the sum of the ECDs predicted for the eight known TM subunits (CATSPER1-4, β, γ, δ, and ε) with 1:1 stoichiometry (Extended Data Table 1). We speculate that each TM auxiliary subunit specifically pairs with a particular pore-forming subunit.

## Intracellular structures of CatSper connect two channel units as diagonal arrays

Markedly, the intracellular domains observed underneath the channel form a continuous diagonal array between two staggered channel complexes of the zigzag string (Fig. 3e, h). The diagonal stripes are spaced by 17.6 nm and are oriented in the same forward slash direction as the two connected inner pore-forming subunits (Fig. 3d). The side view of the complex shows that the intracellular protrusion of an individual channel is not coaxial with the center of the tetrameric channel (Fig. 3a).

The mass estimation of the intracellular domains corresponding to one wild type CatSper channel complex is ∼200 kDa (Fig. 3e, h), which is ∼40 kDa smaller than the combined molecular weights of the cytoplasmic domains from the 10 reported CatSper subunits (Extended Data Table 1). As all the eight known TM subunits are required to make one channel unit^6,21^, these results suggest that two forward slash neighboring channels may form an intercomplex of 2:1 stoichiometry such that an EFCAB9-CATSPERζ pair links two channel units like an intracellular bridge, using two channels as a building block of the zigzag assembly. At this point, it remains unclear whether C2CD6 and TRIM69 would stabilize the CatSper complex or interact rather transiently. Their stoichiometry to other subunits also needs to be determined.

## EFCAB9-CATSPERζ complex has profound impact on the long- and short-range architecture of CatSper channels

Interestingly, we observed that not only the length and alignment of the nanodomain rows was changed in *Efcab9*^-/-^ sperm as compared to wild type sperm (Fig. 2), but also the arrangement between neighboring CatSper complexes were different (Fig. 3i-l and Extended Data Fig. 3c-g). The mutant averages showed that complexes are still arranged in two staggered and anti-parallel lines, as is evident from the preserved location of the wing density (Fig. 3k). However, in the mutant the usual zigzag pattern is disrupted, and instead neighboring complexes forms arrays of diagonal stripes that are oriented either in a backslash (Fig. 3i-k) or a forward slash direction (Extended Data Fig. 3d-f).

In *Efcab9*^-/-^ sperm, the intracellular domain of the CatSper complex is visible but appears reduced (Fig. 3l; Extended Data Fig. 3c, g) and is possibly mis-oriented in the predominant backslash arrangement (Fig. 3l). In the mutant with backslash phenotype, at the roof level the usual forward slash connection is greatly reduced (Fig. 2j, l, 3j) and a different subunit of the tetrameric channel forms the inner connection between neighboring complexes (Fig. 3k). The forward slash arrangement resembles the wild type organization at the tetrameric channel level, but at the roof level the usual backslash connection is missing (Fig. 2n; Extended Data Fig. 3b, e). Despite this re-arrangement, in both configurations, the rows or clusters have a width and ECD mass that is comparable with the wild type zigzag ribbon.

## Similarities and differences between mouse and human sperm CatSper structures

Cryo-tomograms and subtomogram average of human sperm flagella also revealed a ∼24 nm wide zigzag string of staggered and anti-parallel arranged complexes on the extracellular side of the flagellar membrane (Fig. 3m-o). Although the resolution of the subtomogram average of human sperm flagella was limited by a low number of averaged repeats (Extended Data Fig. 3j and Extended Data Table 2), a 11.3 nm wide tetrameric channel with forward slash connection (Fig. 3o), a center-to-center spacing between channels along the zigzag string of 15.2 nm, and the canopy roofs in a zigzag-pattern were clearly visible (Fig. 3n), suggesting that these unique rows (Fig. 2o-r) are likely arrays of human CatSper complexes. We observed only two differences between the mouse and human sperm CatSper complex arrays: first, the human roof ridges were oriented opposite to that of the mouse CatSper, *i*.*e*. in the backslash direction (Fig. 3m), possibly reflecting an ∼45° counterclockwise rotation of each channel unit within the zigzag row; second, human CatSper appears to be missing the wing structure observed for the mouse CatSper tetrameric channel (compare Fig. 3o *vs*. 3d, Extended Data Fig. 5a). The molecular identity of the wing structure remains unclear. However, we propose that this wing consists of SLCO6C1 as it is a rodent-specific, multi-pass TM protein with small ECD, and its quadrilinear localization to the flagellar membrane is dependent on other TM CatSper subunits in mouse sperm (Fig. 1c and Extended Data Fig. 1c).

Physiological substrates are not yet identified for SLCO6C1^17^. However, the International Mouse Phenotyping Consortium reports that *Slco6c1*^-/-^ mice show decreased circulating phosphorus level^20^, suggesting SLCO6C1 could be a potential phosphate transporter. As CatSper is required to sustain motility for extended period of time^7,12^ which in turns is dependent on flagellar energy metabolism^23,24^, the association of SLCO6C1 with murine CatSper channel complex might be a species-specific molecular mechanism linking Ca^2+^ homeostasis to ATP production via glycolysis. Compared with mouse sperm, which use glycolysis as a dominant source of ATP production^25^, human sperm might split ATP production differently between oxidative phosphorylation and glycolysis.

## Structural defects of mutant CatSper correlates with proximally stiff flagellum and compromised motility

The CatSper-mediated increase in intracellular Ca^2+^ initiates in the principal region of the tail and propagates towards the sperm head^26^. Our previous flagellar waveform analyses of head-tethered sperm showed that *Efcab9*^-/-^ sperm display stiff flagella in the proximal region^12^. To better understand how the structural alterations of the mutant CatSper channel complexes are translated into altered flagellar curvature and motility in *Efcab9*^-/-^ sperm, we characterized the flagellar waveform and swim paths of free-swimming sperm in detail over time using 3D high-speed Digital Holographic Microscopy^27^ (Fig. 4, Extended Data Fig. 4). Capacitation dramatically increased *xy*-excursion with respect to the laboratory-fixed frame of reference, i.e. the out-of-plane beating of wild type sperm, which is abolished in *Efcab9*^-/-^ sperm (Fig. 4a, c, d and Extended Data Fig. 4a). By contrast, capacitation did not significantly affect the flagellar *z*-excursion, i.e. the waveform amplitude (Fig. 4a, e), suggesting that Ca^2+^ influx by CatSper activation mainly regulates asymmetric out-of-plane beating in the *xy*-direction, but not flagellar movement in the *z*-direction. Interestingly, the *z*-amplitude of non-capacitated *Efcab9*^-/-^ sperm is smaller than that of wild type sperm (Fig. 4a, e), likely due to the proximally stiff flagellum of *Efcab9*^-/-^ sperm. The proximally stiff mutant flagella might result from a lower basal level of intracellular calcium, balanced by basal CatSper activity and Ca^2+^ extrusion pump^12^.

**Fig. 4.**
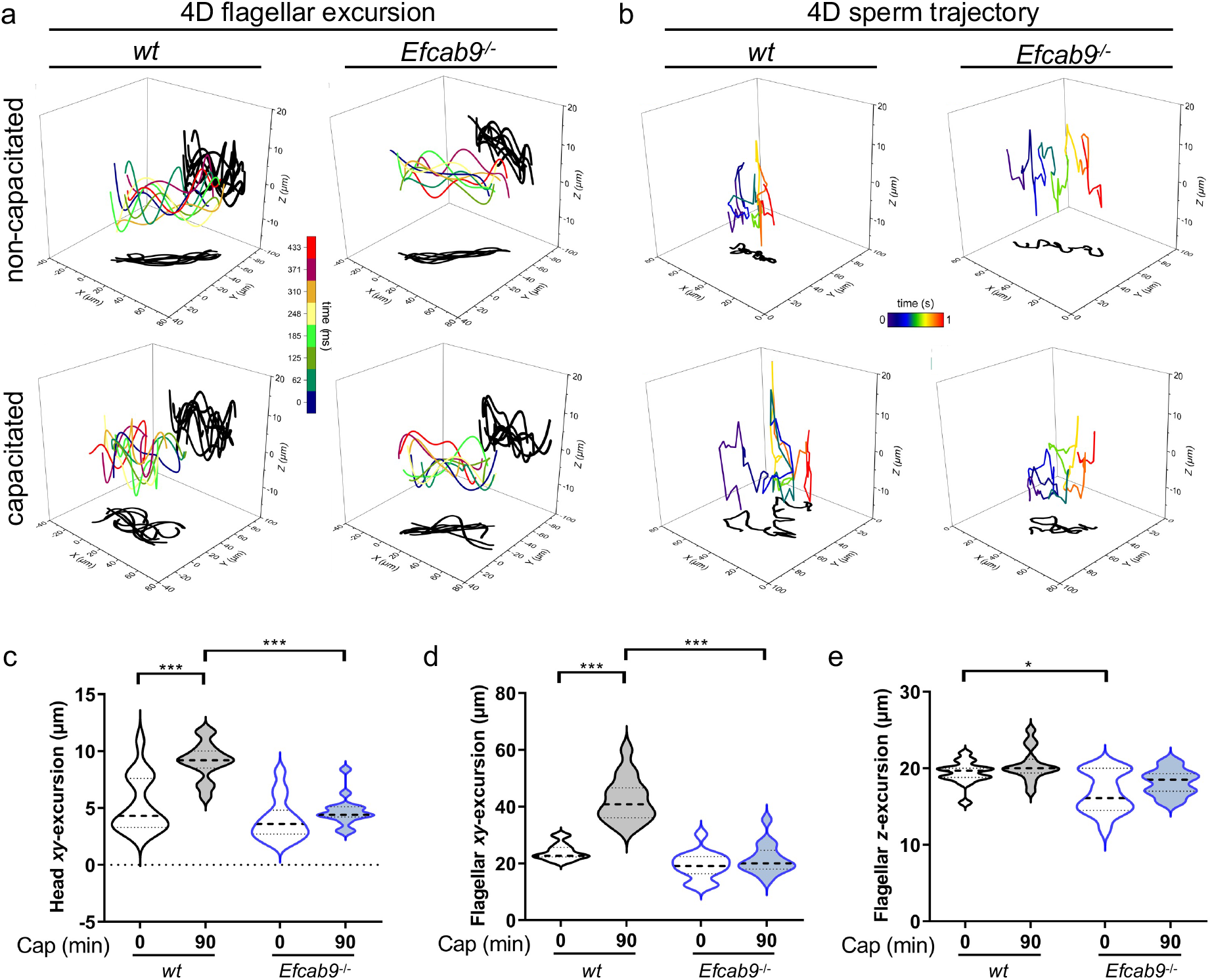
Flagellar beating waveform of free-swimming sperm in four dimensions. **a**. 4D flagellar beating waveform analyses of *wt* and *Efcab9*^-/-^ sperm by high-speed digital holographic microscopy (DHM). The time-lapse trace of a flagellum at 3D position (laboratory-fixed frame of reference *xyz*) is visualized in color and its projections onto *xy*- and *xz*-planes are shadowed in black. **b**. 4D sperm trajectory analyses of *wt* and *Efcab9*^-/-^ sperm by DHM. The swimming trajectory of sperm is visualized by tracing the head position. **c-e**. Statistical analyses of head (**c**) and flagellar (**d**) *xy*-excursion, and flagellar *z*-excursion (**e**) from (**a**). n=15 each group, \**p* < 0.05, \*\*\**p* < 0.001; the medians (thick dash lines) and interquartile ranges (thin dash lines).

To further unravel the effect of the altered beat patterns on sperm swim paths, we determined the 3D trajectories of free-swimming sperm by tracing the head positions using 3D high-speed Digital Holographic Microscopy. Consistent with the increase of curvilinear velocity (Extended Data Fig. 4b), the swimming trajectory of capacitated wild type sperm increase the range of excursion in all dimensions (Fig. 4b, left; compare Extended Data Video. 3 *vs* 4). In contrast, *Efcab9*^-/-^ sperm fail to expand the excursion range during capacitation (Fig. 4b, right), suggesting the importance of CatSper channel and higher-order integrity on the effective sperm navigation during capacitation. Capacitation likely requires Ca^2+^ signaling through the coordinate activity of many CatSper channels. Based on our structural findings and the motility defects in *Efcab9*^-/-^ sperm, we propose that in wild type sperm the extracellular connected zigzag arrangement within a longitudinal nanodomain could coordinate the opening of the entire array of CatSper channels along the flagellar axis, ensuring a large and synchronous Ca^2+^ influx to generate strong bending force (Extended Data Fig. 5b). In contrast, disruption of the CatSper zigzag-rows and misalignment from the longitudinal axis would dysregulate this domino-effect, thus preventing efficient intracellular Ca^2+^ propagation towards the sperm head and resulting in a proximally stiff flagellum and altered sperm motility (Extended Data Fig. 5b). Taken together, this work provides an unprecedented structural basis for understanding the CatSper channel function in motility regulation of mammalian sperm.

## Materials and Methods

### Human subjects

A total of 3 healthy volunteers aged 25-39 were recruited for this study. Freshly ejaculated semen samples were obtained by masturbation and spermatozoa purified by the swim-up technique at 37°C as described in detail in^28^. All processed samples were normozoospermic with a cell count of at least 30 × 10^6^ sperm cells per mL. The experimental procedures utilizing human-derived samples were approved by the Committee on Human Research at the University of California, Berkeley, IRB protocol number 2013-06-5395.

### Animals

*Catsper1*^-/-^, *Catsperd*^-/-^, *Catsperz*^-/-^ and *Efcab9*^-/-^ mice generated in the previous studies^5,8,9,12^ are maintained on a C57/BL6 background. Mice were treated in accordance with guidelines approved by the Yale Animal Care and Use Committees (IACUC).

### Antibodies

Rabbit polyclonal antibodies specific to mouse CATSPER1^5^, δ^9^, and EFCAB9^12^ were described previously. Briefly, to produce antibodies, peptides corresponding to mouse C2CD6 (ALS2CR11) (359-377, EKLREKPRERLERMKEEYK) (Open Biosystems) and SLCO6C1 (1-14, MAHVRNKKSDDKKA) (GenScript) were synthesized and conjugated to KLH carrier protein. Antisera from the immunized rabbits were affinity-purified using the peptide immobilized on Amino Link Plus resin (Pierce). Other antibodies used in this study are commercially available as follows (TRIM69, Origene; Fancm, Affinity Biosciences; acetylated tubulin, Sigma). All the chemicals were from Sigma Aldrich unless otherwise indicated.

### Western blot analysis

Whole mouse sperm protein content was extracted as previously described^8,9,14^. In short, mouse epididymal spermatozoa washed in PBS were directly lysed in a 2×SDS sample buffer. The whole sperm lysate was centrifuged at 15,000 g, 4°C for 10 min. After adjusting DTT to 50 mM, the supernatant was denatured at 95°C for 10 min before loading to gel. Antibodies used for Western blotting were antibodies against CATSPER1 (1 µg/mL), δ (1 µg/mL), EFCAB9 (1 µg/mL) and C2CD6 (1 µg/mL), TRIM69 (0.5 µg/mL), SLCO6C1 (2 µg/mL), FANCM (1 µg/mL) and acetylated tubulin (1:10000 µg/mL). Secondary antibodies were anti-rabbit IgG-HRP (1:10,000), anti-goat IgG-HRP (1:10,000) and anti-mouse IgG-HRP (1:10,000) from Jackson ImmunoResearch (West Grove).

### Sperm immunocytochemistry

Sperm were washed in PBS twice, attached on glass coverslips, and fixed with 4% paraformaldehyde (PFA) in PBS at room temperature (RT) for 10 minutes (mouse) or at 4°C for 1 hr (human). Fixed samples were permeabilized using 0.1% Triton X-100 in PBS at RT for 10 minutes, washed in PBS, and blocked with 10% goat serum in PBS at RT for 1 hr. Cells were stained with anti-C2CD6 (10 μg/mL), TRIM69 (5 μg/mL), SLCO6C1 (10 μg/mL), FANCM (10 μg/mL) in PBS supplemented with 10% donkey serum at 4°C overnight. After washing in PBS, the samples were incubated with donkey anti-goat Alexa 568 (Invitrogen, 1:1,000) in 10% donkey serum in PBS at RT for 1 hr. Hoechst was used to counterstain nuclei for sperm head visualization. Immunostained samples were mounted with Prolong gold (Invitrogen) and cured for 24 hr.

### Confocal and 3D structured illumination microscopy (SIM) imaging

Confocal imaging was performed on the Cured samples by a Zeiss LSM710 using a Plan-Apochrombat 63X/1.40 and an alpha Plan-APO 100X/1.46 oil objective lens (Carl Zeiss). 3D SIM imaging was performed with a Zeiss LSM710 Elyra P1 using an alpha Plan-APO 100X/1.46 oil objective lens. A laser at 561 nm (200 mW) was used for Alexa 568 (Invitrogen). A z-stack was acquired from 42 optical sections with a 200 nm interval. Each section was imaged using 5 rotations with a 51 nm grating period. 3D SIM Images were rendered using Zen 2012 SP2 software.

### Sperm sample preparation for cryo-electron microscopy

Epididymal spermatozoa from adult male mice (wild type and *Efcab9*^-/-^ in the C57BL/6 background) were collected by swim-out from caudal epididymis as described^29^. Briefly, male mice were euthanized, and the cauda isolated from the mouse carcass and placed into a 1.5 mL tube with standard HEPES saline HS medium (in mM: 135 NaCl, 5 KCl, 1 MgSO_4_, 2 CaCl_2_, 20 HEPES, 5 D-glucose, 10 Lactic acid, 1 Na pyruvate, pH 7.4 adjusted with NaOH, osmolarity 320 mOsm/L) at room temperature. To retrieve the mature spermatozoa, the caudal epididymis was cut into several pieces with a scalpel and placed into a 37°C incubator for 10-30 min to let the sperm swim out into the HS buffer. Then, the sample was placed at room temperature for 30 min to let the debris sediment by passive sedimentation, before separating the supernatant with swimming sperm cells from the debris. The supernatant with the sperm was washed one time in PBS, which involved centrifugation at 700g for 5 min at room temperature.

Human sperm were allowed to settle at the base of a conical tube and the excess buffer was removed. The sperm sample was then passed three times through a Balch ball bearing homogenizer (Isobiotech, 15μm clearance).

### Cryo-sample preparation for cryo-ET

Small aliquots of freshly prepared mouse sperm at a concentration of 1-5 × 10^6^ cells/mL were gently mixed with 10-fold concentrated, BSA-coated 10-nm colloidal gold solution (Sigma Aldrich) at 3:1 ratio, before applying 4 µL of the solution to a glow-discharged (30s at 35 mA) copper R 2/2 200-mesh holey carbon grid (Quantifoil Micro Tools). The grids were blotted manually from the back side with Whatman filter paper #1 for 2-4 s, before plunge-freezing the grid in liquid ethane using a homemade plunge-freezer. Grids were stored under liquid nitrogen until either further preparation by cryo-focused ion beam (FIB) milling or imaging by cryo-ET. For mechanical support, grids were mounted into Autogrids (ThermoFisher).

3 μl of the human sperm sample were applied to glow discharged copper R2/2 200-mesh holey carbon grid (Quantifoil Micro Tools) and plunge frozen in liquid ethane using an automatic plunge freezer (Vitrobot, FEI, blot force 8, blot time 8s, Whatman filter paper #1).

### Cryo-electron tomography

Tilt series of whole or cryo-FIB milled mouse sperm flagella were acquired using a Titan Krios (Thermo Fisher Scientific) operated at 300 keV with post-column energy filter (Gatan) in zero-loss mode with 20 eV slit width. Images were recorded using a K3 Summit direct electron detector (Gatan) in counting mode with dose-fractionation (12 frames, 0.05s exposure time per frame, dose rate of 28 electrons/pixel/s for each tilt image). Tilt series were collected using SerialEM^30^ with the Volta Phase Plate and a target defocus of -0.5 μm. Images were recorded at 26k × magnification resulting in a pixel size of 3.15 Å. Dose-symmetric tilt series^31^ were recorded under low-dose conditions, ranging from ±60° with 2° angular intervals with the total electron dose limited to ∼100 e^-^/Å^2^.

Frozen grids of human sperm were loaded into a Jeol3100 TEM operating at 300kV equipped with an in-column energy filter and a direct electron detector (K2, Gatan). Dose-fractionated, bi-directional tilt series were acquired using SerialEM^30^ with the following parameters: angular increment 1.5°, angular range about +/-60° starting at -20°, energy filter slit width 30 eV, nominal magnification 10k × resulting in a detector pixel size of 3.98 Å (which was binned by x2 resulting in a pixel size of 7.96 Å in the reconstruction), defocus -2.5 μm, exposure time 1s × 1/cos(tilt angle), fraction interval 0.2 s, dose rate 1 e^-^/Å^2^/s, total dose ∼80 e^-^/Å^2^.

### Cryo-FIB milling of mouse sperm

For lamella (section) prepared by cryo-FIB-milling, clipped grids (modified Autogrids with FIB notch) with plunge-frozen mouse sperm were transferred to an Aquilos dual-beam instrument with cryo-sample stage (Thermo Fisher Scientific). Two layers of platinum were added to the sample surface to enhance sample protection and conductivity (sputter-coater: 1 keV and 30 mA for 20s; gas injection system (GIS): when needed, heated up to 28°C, and then deposited onto the grid for 15 seconds)^32^. Scanning electron beam imaging was performed at 2 kV and 25 pA, and Gallium ion beam imaging for targeting was performed at 30 kV and 1.5 pA. The target region, *i*.*e*. a sperm flagellum, was oriented for milling by tilting the cryo-stage to a shallow angle of 14 - 16° between the ion beam and the grid. Cryo-FIB milling was performed using a 30 keV gallium ion beam with a current of 30 pA for bulk milling, 30 pA for thinning, and 10 pA for final polishing, resulting in 100-200 nm thick self-supporting lamella, that could then be imaged by cryo-ET.

### Image processing of cryo-ET data

For tilt series of both mouse and human sperm flagella, movie frames were aligned using Motioncor2 1.2.3^33^. The IMOD software^34^ was used to align the tilt serial images using the 10-nm gold particles as fiducial markers and to reconstruct the tomograms by weighted back-projection. For subtomogram averaging, the repeating units were picked manually from raw tomograms. The repeat orientation was determined based on the polarity of the axoneme at the core of the sperm flagella. The alignment and missing-wedge-compensated averaging were performed using the PEET software^35^. After initial averaging a two-fold symmetry was applied. Visualization of the 3D structures of the averaged repeat units was done using the UCSF Chimera software package^36^. Mass estimations from a repeat unit in the subtomogram averages were calculated using the average density of 1.43 g/cm^3^ for proteins^37^ and normalization of the isosurface-rendering threshold in Chimera. The number of tomograms of whole cell and lamella, number of averaged repeats and estimated resolutions of the averages (using the FSC 0.5 criterion), are summarized in Extended Data Table 2.

### Analysis of mouse sperm motility and flagellar beating in 4D

Epididymal spermatozoa from adult male mice were collected by swim-out from caudal epididymis in standard HEPES saline HS medium.

For 4D analysis, mouse sperm were washed twice in HS medium and resuspended to a final concentration of 1-2 ⨯ 10^6^ cells/mL either under non-capacitating (HS medium) or under capacitating (HS medium, 15 mM NaHCO_3_^-^, 5 mg/mL BSA) conditions. To induce capacitation *in vitro*, sperm were incubated for 90 min at 37°C and 5% CO_2_. 4D motility analysis was done at 37°C and 5% CO_2_ using an off-axis transmission digital holographic microscope DHM™ T-1000 (Lyncée Tec SA, Geneva, Switzerland) equipped with a 666 nm laser diode source, a 20x/0.4 NA objective and a Basler aca1920-155um CCD camera (Basler AG, Ahrensburg, Germany). Holographic imaging was performed as previously described^27^. In short, mouse sperm were placed in a 100 µm deep chamber slide (Leja) and were recorded at 100 Fps. Offline processing was done using proprietary Koala (Vers. 6; Lyncée Tec SA) and open-source Spyder (Python 3.6.9) software. Using Koala software, *xy*-plane (parallel to the objective slide) projection images of sperm were numerically calculated at different focal planes (z-height) ^38,39^, followed by sperm head tracking using Spyder to receive *x, y* and *z*-coordinates for the entire trajectory. Using these coordinates, motility parameters including 3D curvilinear velocity (VCL, in μm/s) and 2D amplitude of lateral head displacement (ALH, in μm) were analyzed. For each condition, 15 free-swimming single sperm were analyzed using three males from each genotype (wild type, *Efcab9*^*-/-*^, *Catsper1*^*-/-*^).

For 4D flagellar beating analysis, a macro written in Igor Pro™ Vers. 6.36 (Wavemetrics) was used to perform frame-by-frame tracking of flagellar images in stacks of reconstructed *xy*-projections (8-bit TIFF format, 100 Fps, 10 Frame Time) with a resolution of 800 × 800 pixels as well as automatic brightness and contrast adjustments applied by ImageJ V1.50i (National Institutes of Health). A P/U value (3.7466), which is defined as the quotient from the objective magnification (20x) and the pixel size (5.34 μm) of the camera (Basler aca1920-155um), was used to convert pixel to micrometer. Calculation of *z*-coordinates was performed utilizing the received *x, y*-coordinates from flagellar traces and Koala. A specific script in Spyder was used to load flagellar *x, y*-coordinates into Koala. Smoothing of *z*-plane data was conducted with Igor Pro™ by fitting to 7^th^ order polynomials. The determination of the distance along the flagellum in the *xy* projections (Dx, y) was carried out geometrically from adjacent pairs of *x, y-*coordinates, also using a macro in Igor Pro™.

4D visualization of sperm flagellum and sperm swimming trajectories with respect to the laboratory fixed frame of reference (*x, y, z*) was done using OriginPro 2020 software (OriginLab Corporation). Therefore *x, y*, and *z*-coordinates of head and flagellar tracking were imported to the software. Analysis was performed for one whole beat-cycle, but for better illustration only, every 6^th^ flagellar excursion between maxima of one beat cycle was illustrated (frame 0, 6, …, 42, every 60 ms) in Fig. 4. The associated movies (Extended Data Videos 3 and 4) of reconstructed trajectories of free-swimming single sperm were created with Cinema4D Vers. 18 (Maxon) using *x, y*, and *z*-values of the 4D head tracking. Adobe After Effects software Vers. CS6 (Adobe Systems Software Ireland Limited) was used for video composing and time duration adding. In each supporting video two different perspectives were used to show the 3D movement of sperm during 1s record. The rolling ball represents the sperm head, and the color code of the trajectory displays the *z*-excursion.

### Quantification and statistical Analysis

Statistical analyses were carried out with GraphPad Prism 9 (Statcon GmbH) by using a two-way analysis of variance (ANOVA). Differences were considered significant at *p* <0.05. Numerical results are presented as medians and interquartile ranges with n = number of determinations and N = number of independent experiments.

## Data and software availability

The averaged 3D structure of CatSper channel from mouse wild type sperm flagella has been deposited in the Electron Microscopy Data Bank (EMDB) under accession code EMD-24210.

## Acknowledgements

Authors thank Drs. Jun Liu and Shiwei Zhu for help in initial trials of mouse sperm cryo-ET at the Yale West campus Cryo-EM core, Dr. Daniel Stoddard for providing EM training and management of the UT Southwestern Medical Center (UTSW) cryo-electron microscope facility (funded in part by a Cancer Prevention and Research Institute of Texas Core Facility Award (RP170644)), David Mastronarde and John Heumann (University of Colorado at Boulder) for continued development of image processing tools, Jürgen Heger for the trajectory videos, the BioHPC supercomputing facility located in the Lyda Hill Department of Bioinformatics at UTSW for the computational resources and Dr. Fred Sigworth for critically reading of the initial draft. This work was funded by the National Institutes of Health (R01HD096745 to J.-J.C, R01GM083122 to D.N, R01GM111802 to P.V.L.), the Cancer Prevention and Research Institute of Texas (RR140082 to D.N.), the Grantham Foundation to J.-J.C. the Pew Biomedical Scholars Award and Rose Hill award to P.V.L., in part by the Office of Science of the US Department of Energy (DE-AC02-O5CH11231) and UCB start-up funds to K.M.D., and the Deutsche Forschungsgemeinschaft (2344/9-3 to G.W.). J.Y.H. and X.H. are recipients of the Male Contraceptive Initiative and James Hudson Brown-Alexander B. Coxe Postdoctoral Fellowship, respectively.

## Author Contributions

J.-J.C. and P.L. conceived the study. D.N., and J.-J.C. designed and oversaw the project. H.W., J.Y.H., and X.H. performed biochemical characterization of novel CatSper components and confocal and SR imaging experiments. Y.Z., H.W., and N.B.S. performed EM sample preparation and screening. Y.Z., D.N. (mouse sperm) and N.B.S., K.M.D. (human sperm) performed cryo-ET, Y.Z. performed subtomogram averaging and 3D visualization. E.R. performed cryo-FIB milling. C.W. performed sperm motility experiments and 4D flagellar beating analysis. Y.Z., H.W., and C.W. made figures. Y.Z., H.W., P.L., K.M.D., G.W., D.N., and J.-J.C. interpreted data. H.W., and J.-J.C. prepared the initial draft of the manuscript. Y.Z., H.W., C.W., G.W., P.L., K.M.D., D.N, and J.-J.C. edited the manuscript with input from all other authors in the final version. P.L., K.M.D., G.W., D.N., and J.-J.C. obtained funding.

